# Bi-clustering interpretation and prediction of correlation between gene expression and protein abundance

**DOI:** 10.1101/270397

**Authors:** Xiaojun Wang, Lin Teng, Haicang Zhang, Qian Zhou, Xiaoquan Su, Xinping Cui, Dongbo Bu, Xinqi Gong, Ansgar Poetsch, Kang Ning

**Author notes:** Equal contributors (Xiaojun Wang, Lin Teng). Corresponding author: Kang Ning Ansgar Poetsch Xinqi Gong.

## Abstract

Most organisms’ transcript and protein level only moderately correlate for various reasons, such as regulation of transcription and protein degradation. Better prediction and understanding the correlation between gene expression and protein abundance has been possible by harnessing the matching RNA/protein datasets produced by modern high-throughput RNA-Seq and mass spectrometry methods. In this work, we have utilized some well-studied matching RNA/protein datasets, and explored for the first time a bi-clustering method to cluster genes that have consistent correlation patterns between gene expression and protein abundance. The clustering results have been interpreted from the perspective of both transcriptomic and proteomic features, which show that mRNA half-life, protein half-life and protein structure in concert significantly affect the correlation of gene expression and protein abundance. With these and other carefully selected features, a prediction model based on individual clusters, called Cluster-based Linear prediction Model (CLM), was built and tested on mouse liver mitochondrial, mouse brainstem mitochondrial, *Saccharomyces cerevisiae* and *Danio rerio* datasets. CLM could find genes for which protein abundance can be predicted from mRNA data. In summary, based on bi-clustering, feature selection and CLM model, we have established a new and valuable cluster-based protein abundance prediction method.

## Introduction

In recent years, it has become obvious that the amounts of matching mRNAs and proteins in a cell are not only governed by mRNA and protein synthesis, but also by intricate regulatory processes, such as regulatory RNA and protein modifications^1^. Hence, based on studies with many species, the general conclusion is that cellular mRNA and protein levels only correlates weakly^2–9^: the squared Pearson’s correlation coefficient(R^2^) is ~0.4 for gene expression and protein abundance, implying that only ~40% of the variance in protein abundance can be explained by changes at the transcript level^10^. And analyses by *Ning* et al.^11^ comparing RNA-Seq and mass-spectrometry-based protein abundance data also led to a similar conclusion of modest correlation. Several other investigations reported slightly higher correlation, but these were limited to a few hundred mRNA-protein pairs (0.7)^12^, or differential expressed genes (0.88)^13^. Thus, integration of proteomic and transcriptomic approaches is of great importance, especially for biomarker identification in disease diagnosis^14^. Admittedly, quantitative proteome analysis with mass spectrometry has undergone tremendous improvements in the last decade^13,15,16^, yet it still cannot compete in terms of coverage, sensitivity and dynamic range with RNA-Seq. Due to these technical limitations, we are facing the dilemma that the biologically often more relevant description of protein abundance is unachievable and adequate approaches are still required to predict protein abundance based on gene expression data and additional features.

Whereas it remains impossible to account for all regulatory principles when predicting protein abundance from gene expression, some have been implemented and essentially three kinds of approaches can be distinguished: (1) approaches based on models considering protein synthesis and degradation; (2) approaches based on models simulating the protein translation process; (3) approaches utilizing various gene properties such as mRNA and protein sequences and 3D structures.

Regarding the first kind of approaches, many groups tried to develop mathematical models to predict the changes of protein expressions from translation rate of mRNA and degradation rate of protein^13,17,18^. Although their efforts faced many obstacles, such as technical difficulties in measuring these two rates and applicability of the model only to cells in steady state^19^, such mathematical model should have significant implications in research on translation or degradation regulation. For the second, utilizing models based on simulating the protein translation process, some groups have built models to predict key translation rates (i.e., Ribosome Flow Model based on Totally Asymmetric Exclusion Process^20^) in the translation process. Based on mathematical models, prediction of protein abundance from gene expression would become possible. However, protein abundance levels had to be determined by balancing protein production and degradation rates - which are hardly available.

For the approaches that utilize various gene properties, researchers have previously utilized all-gene-based General Linear Model (GLM) to correlate gene expression with protein abundance^11^, but little biological insight was gained except for global correlation. Some groups tried to improve protein abundance prediction by building models that combined protein or mRNA features such as sequence frequencies and properties with gene expression^21,22^. One of these is the Multivariate Adaptive Regression Splines (MARS) model^23^, in which each function is constructed to fit distinct intervals of variables, and in each function piecewise linear segments (referred to as splines) are smoothly connected. MARS model separates all genes into clusters, and for each cluster a linear relationship could be obtained. Such linear relationship would always result in better values (than global linear relationship value) since these clusters are generated to optimize the inner consistency of genes/proteins. Thus “divide and conquer” strategies would usually work better than a GLM (or similar) strategy. However, as MARS was modeled as a combination of piecewise continuous linear functions, the splines in MARS model were built without considering any biological mechanism, making biological interpretation of splines difficult.

In this work, we utilized some well-studied matching RNA/protein datasets^11^ to explore the dynamic relationship between transcript and protein expression, for which: (1) We have evaluated the mouse liver mitochondrial dataset, and selected representatives from 18 methods that were most suitable for protein, and 2 methods most suitable for mRNA abundance calculation^3^. Gene groups with consistent proteome and transcriptome data were discovered with hierarchical clustering through the QUBIC algorithm^24^ (details in **Supplementary Information**). (2) We have interpreted the bi-clustering results mainly from three different aspects, including mRNA half-life, protein half-life and protein 3D structure, to identify the general features of gene expression products in each subgroup after bi-clustering. (3) We have also quantified the general characters of gene products (protein sequence length, etc.) from the bi-clustering results, and used them to develop a mathematical prediction model (CLM, a “divide and conquer” strategy) that could be used to predict protein abundances from corresponding gene expression. (4) Finally, the prediction method (CLM modeling approach) was validated on the mouse brainstem mitochondrial dataset, a *Saccharomyces cerevisiae* dataset, as well as a *Danio rerio* dataset, based on which the advantage of our proposed framework was confirmed effective for feature selection as well as prediction model building.

## Materials and Methods

### Datasets

Currently more and more corresponding mRNA/protein datasets are becoming available^11,13,25^. In this work, we selected four different sets of High-Throughput Sequencing (HTS) transcriptome data and mass spectrometry data (normalized) were used to analyze the relationship between gene expression and protein abundance, as well as properties of genes for: 1. mouse liver mitochondria^11^, 2. mouse brainstem mitochondria^11^, 3. *Saccharomyces cerevisiae^25^,* and 4. *Danio rerio*^26^. We have used mouse liver mitochondria dataset as the main dataset for building and testing the CLM modeling approach. This dataset contained 442 genes in total (after biclustering), and each sample had the corresponding gene expression and protein abundance values determined by different methods, with half-lives of protein and mRNA derived from Schwanhausser *et al.*^17^. Additionally, the other three datasets were used for validation of our approach: (1) The mouse brainstem dataset has similar data structure (380 genes with both corresponding gene expression and protein abundance values) with the liver dataset, differing only in tissue sources. (2) *Saccharomyces cerevisiae* dataset containing isotope-coded affinity tag (ICAT) for protein abundances and microarray methods for gene expressions. (3) *Danio rerio* embryonic development dataset (5001 genes in total) containing NanoHPLC-ESI-MS/MS data for protein abundances and RNA-Seq methods from gene expressions. These three datasets were used to validate the applicability of our approaches in different species. The use of genes from mouse brainstem tissue, as well as genes in *Saccharomyces cerevisiae* and *Danio rerio* would validate the applicability of our analytical approaches and prediction models for establishment of the connection between gene expression and protein abundance.

**Figure 1.**
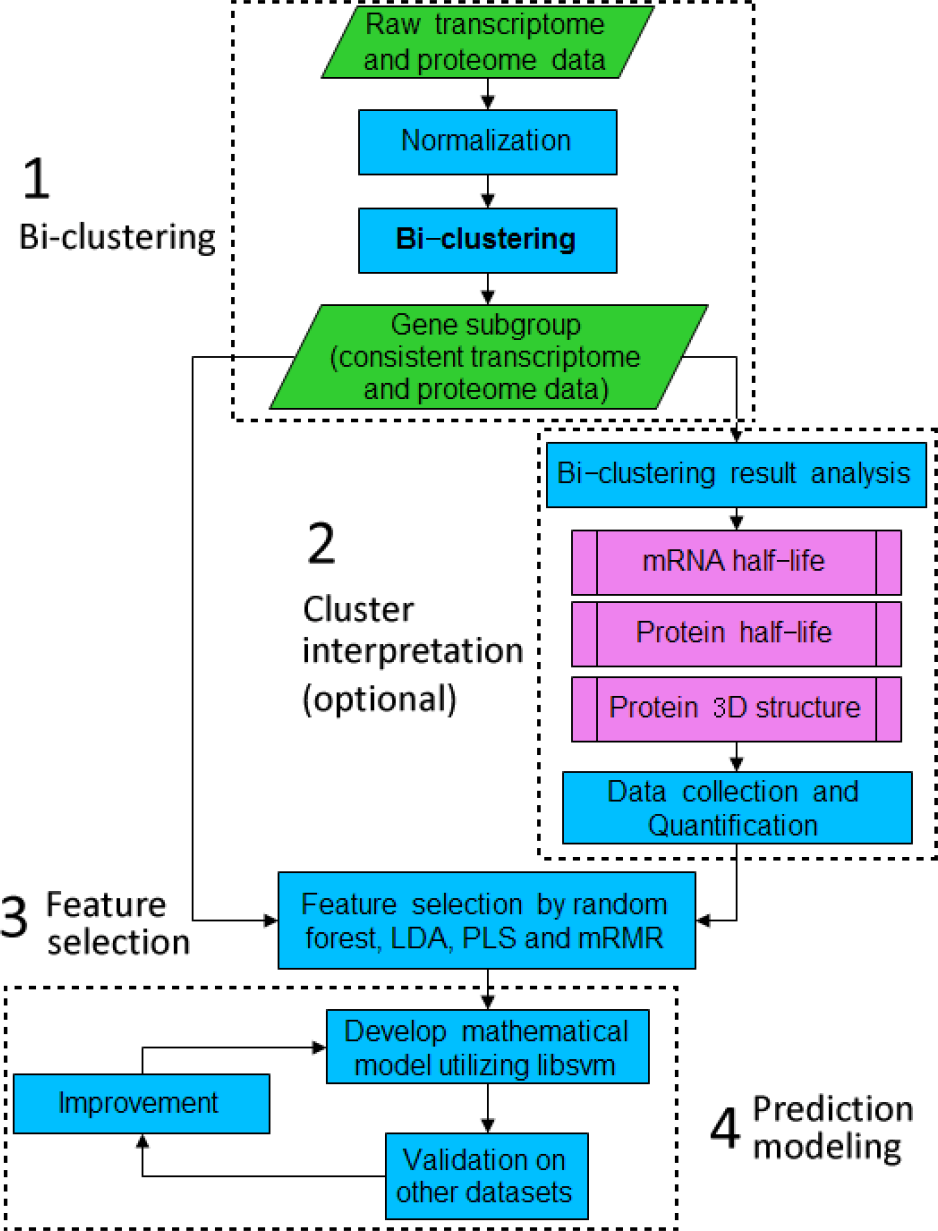
The workflow for bi-clustering, cluster interpretation and prediction modeling. The whole process includes 4 major components: (1) bi-clustering, (2) cluster interpretation (optional), (3) feature selection, (4) prediction modeling.

### Prediction workflow

The overall prediction workflow was depicted in **Figure 1**.

#### (1) Gene bi-clustering and characterization

Suitable calculations for gene expression and protein abundance were chosen before gene clustering. For protein abundance data, we could select from 18 different calculation methods^11^ (**Figure S1**). However, since some of these methods are highly correlated in detecting protein abundance, we performed hierarchical clustering by IBM SPSS’s modules on these methods and selected one method from each cluster to avoid redundancy. 4 methods were kept thereafter. Based on these 4 methods, together with 2 methods for gene expression analyses, the gene expression and protein abundance measurement for mouse liver mitochondria genes were used as input (a matrix) for bi-clustering. Our goal was that (1) clusters contain at least 15 genes and (2) the gene/protein expression trend for genes in these clusters are consistent as judged by at least 1 protein abundance and 1 gene expression method, and we adjusted the parameters (*q* = 0.5 and *r* = 2) accordingly.

#### (2) Cluster interpretation

We interpreted the relationship of gene expression and protein abundance in each one of the obtained clusters from different aspects. For mRNA, the factors that can influence its half-life or stability (e.g., RNA secondary structure) varied. For protein, we considered its function and 3D structure (considering the complexity of protein 3D structures, only the ratio of “protein surface area/volume” was calculated using online server (SARpred, http://www.imtech.res.in/raghava/sarpred/)^27,28^ and observed that the higher the ratio, the more prone the protein to degrade), as well as other factors such as protein length (**Table 1**).

**Table 1.**
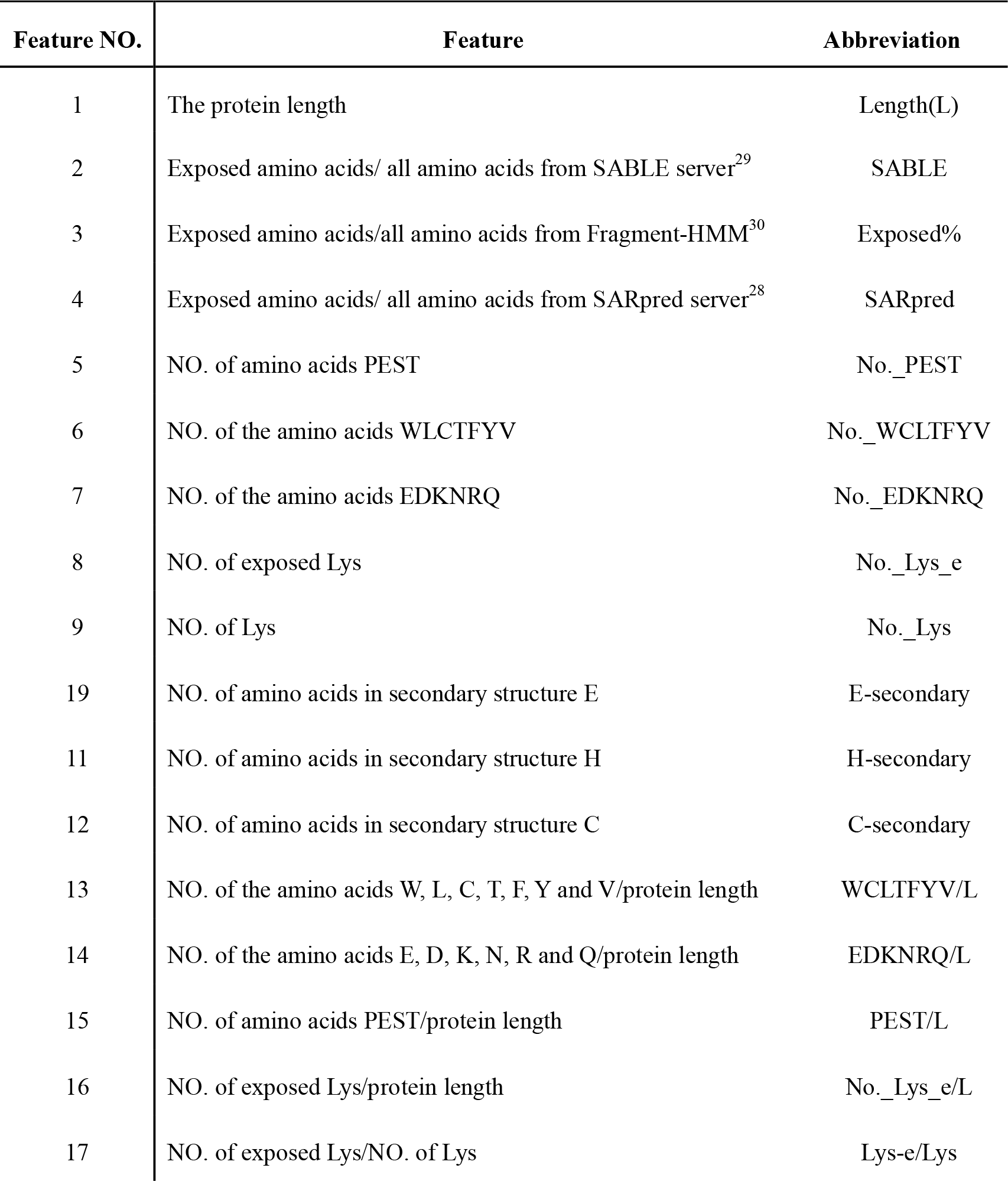
The list of features considered and their abbreviations.

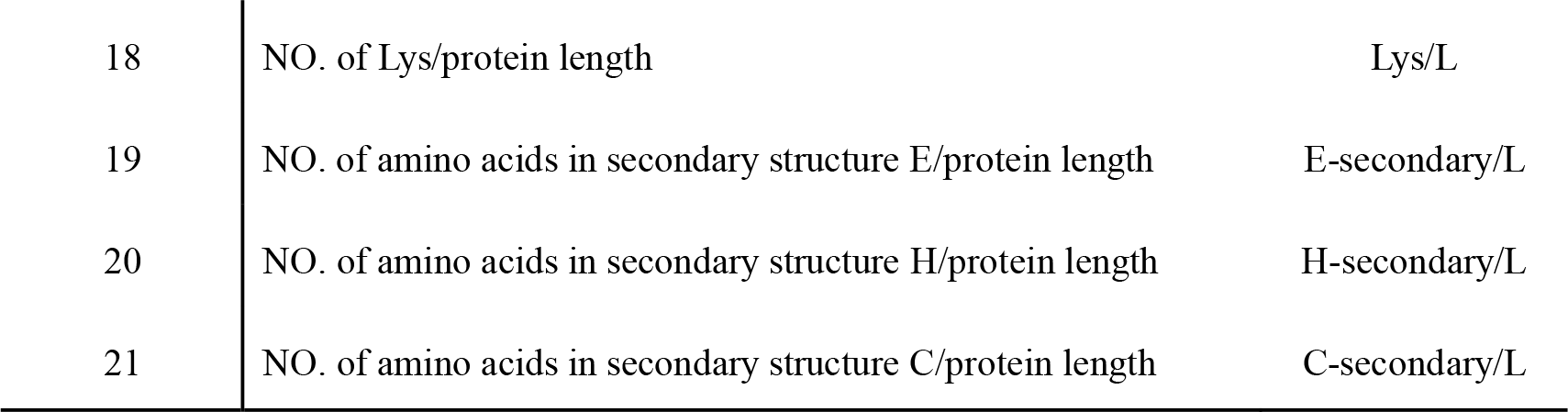

#### (3) Feature selection

Based on gene bi-clustering and characterization, we have observed that many mRNA or protein features were related to mRNA and protein half-lives, thus affecting clustering results. Therefore, 21 important features that related to protein sequences and protein 3D structures were selected according to literatures^31–34^ (listed in **Table 1**). These features would be useful for building the prediction models, based on which gene expression and protein abundance could be connected. 4 different feature selection methods were employed for obtaining a minimal set of discriminatory features: Random forest, PCA (Principal Component Analysis) and LDA (Linear Discriminative Analysis), PLS (Partial least squares regression), and mRMR^35^ (minimum Redundancy Maximum Relevance Feature selection) (more details in **Supplementary Information**).

#### (4) Prediction model

The prediction model that we have proposed and compared here referred to actual gene expression and protein abundance correlation models, based on which prediction of protein abundance from gene expression could be realized. In addition, the selected features for genes in each cluster had to be quantitative (e.g. sequence length or other properties), and we applied them to develop the prediction model that can accurately predict the correlation between gene expression and protein abundance. *Classification and prediction*: The Libsvm^36^ classification method was employed to conduct classification and prediction of gene categories (100 times for each process), then Cluster-based Linear Model” (CLM) for prediction of protein abundance from gene expression was built in each cluster. Thereafter, the CLM prediction model was compared with GLM and MARS model in this study, using the whole genes in mouse liver mitochondrial dataset as the input data. *Accuracy determination:* all the predicted genes were firstly randomly divided into 7 parts; secondly, each part of genes was predicted by the linear model based on genes in the corresponding cluster; thirdly, the prediction process for each model was done many times to obtain an average accuracy result (**Figure S2**); finally, the average accuracy based on the 7 clusters was our prediction result on the total dataset.

## Results and Discussions

In this work, we first established the bi-clustering method, then performed feature selection, and developed the prediction model with -omics data from mouse liver mitochondria. Thereafter, our new workflow was validated with datasets from mouse brainstem mitochondria, *S. cerevisiae* and *Danio rerio.*

### Bi-clustering results and interpretation of clusters

#### (1) Selection of quantification methods

First, we have performed bi-clustering analysis based on the mouse liver mitochondrial dataset. To calculate protein abundance, we initially selected NSAF (MS2 based: using spectrum counting information from MS/MS experiment results), msInspect (MS1 based: using peak intensity information from MS experiment results), msBID (MS1 based) and SpecturmMill (MS1 based) methods. These commonly used methods are selected because NSAF is based on spectral count, whereas the other three methods are based on MS signal intensity, yet use different algorithms^11^. As each of msInspect and msBID has 8 configurations, we could choose from overall 18 different measurements for protein abundances. To avoid bias in quantification methods /algorithms, guided by hierarchical clustering (the agglomeration method=“complete”) result (**Figure S1**), msInspectTop3absPeakIntensity, msBIDTop3normAreaIntensity, SpectrumMill and NSAF were chosen. Based on these 4 methods, together with 2 methods for mRNA analyses (microarray and RNA-Seq), the gene expression and protein abundance measurements for mouse liver mitochondria genes were normalized, and these normalized expression values were used as input for bi-clustering (due to the strict filtration, only hundreds of genes were kept as the input data).

#### (2) Bi-clustering of genes

By using bi-clustering algorithm QUBIC, we have obtained 7 clusters (**Figure 2** and **Figure S3**), each of which had more than 15 genes. Out of these clusters, 4 main clusters each having relatively large number of genes and also having clear half-life patterns were shown in **Figure 3**. These clusters were selected by bi-clustering algorithm, which would have reflected intrinsic properties of gene expression that might not be understood *a priori*.

**Figure 2.**
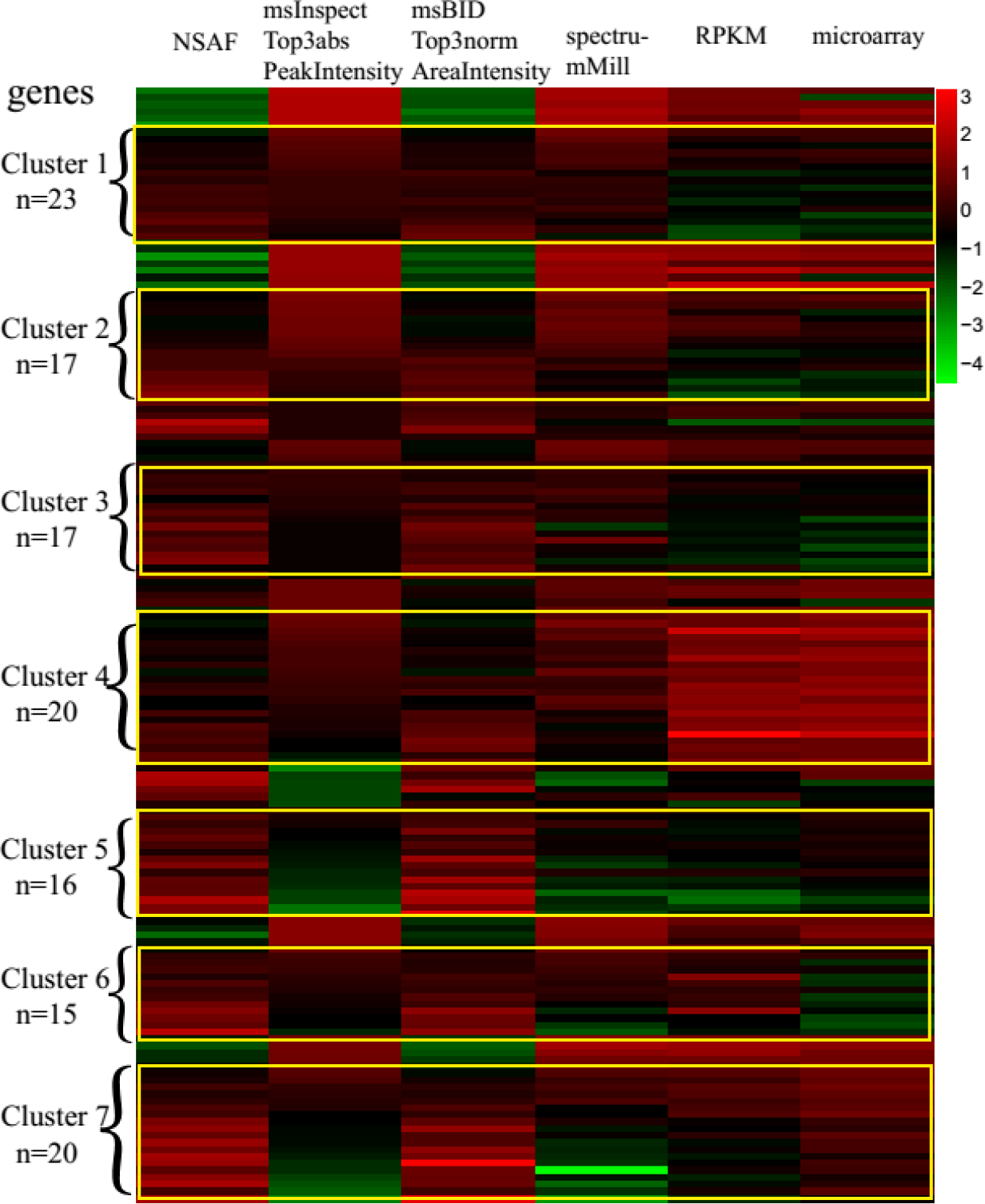
Bi-clustering results for mouse liver mitochondrial genes. The results of bi-clustering were presented in a matrix (442 genes * 6 methods). We have obtained 7 clusters and each cluster contained different number of genes. Values shown in heatmap represented the normalized measurements by different methods.

Despite the weak correlation between gene expression and protein abundance, the joint analysis of transcriptome and proteome data taken from the same samples can potentially reveal clusters of (a) genes with both stable mRNA and stable protein, such as Cluster 1; (b) genes with stable mRNA and unstable proteins, such as Cluster 3 and 4; (c) genes with stable proteins and unstable mRNA, such as Cluster 2; and (d) genes with both unstable mRNA and unstable proteins (**Table S1**). The analysis of these gene clusters in respect to mRNA half-life, protein half-life, protein 3D structure and other features, might further unfold the underlying dynamic relationship between gene expression and protein abundance. The half-life properties and protein functions for four representative clusters were explained as below:

**Figure 3.**
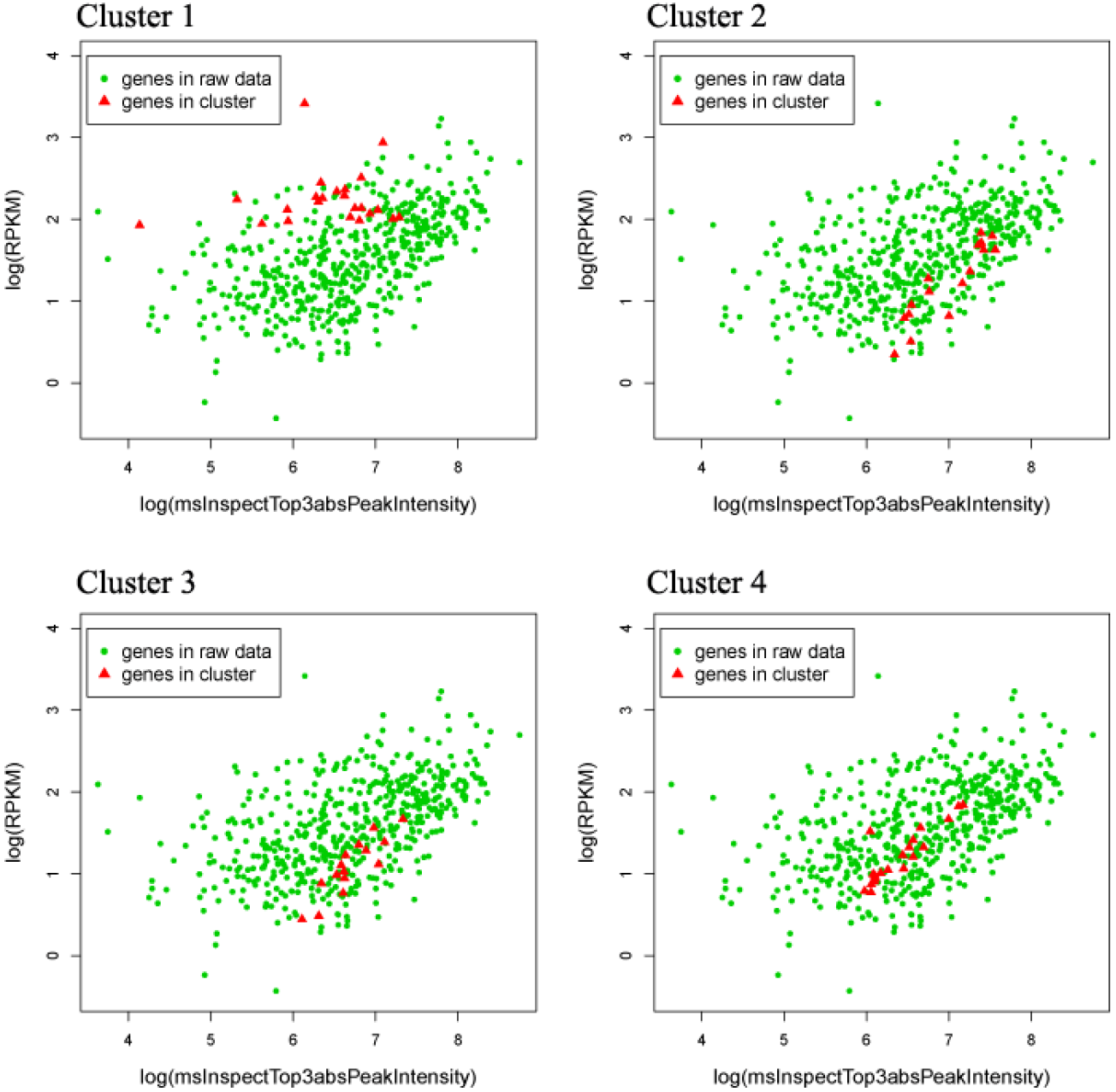
Clusters after bi-clustering for mouse liver mitochondrial genes. The scatter plots of gene distributions for 4 clusters. The red points indicate genes contained in each cluster, respectively. The horizontal axis is the log2 value of protein abundances determined by the msInspectTop3absPeakIntensity method. Vertical axis is the log2 value of gene expression (RPKM values).

##### (i) Cluster 1

23 genes were classified in this cluster. The distribution of gene expression products for the whole population was shown in **Figure 3** (Cluster 1). We observed that gene expressions in log (RPKM) were between 2 and 2.5. These gene expressions were consistent, but the distribution of corresponding protein abundance was dispersed. We conjectured that the stability of mRNA might account for their being in the same cluster. This seemed reasonable, as the average half-life of all mRNA was 12.06 hours, while the average half-life of mRNA in this cluster (15.76 hours) was significantly longer according to *t-test* (*p-value* < 0.001) (**Figure 4a**). It was clear that the genes in cluster 1 can be divided into two sub-groups based on protein half-lives: one included some housekeeping genes with stable mRNAs and stable proteins, such as *HAGH* (encodes hydroxyacyl glutathione hydrolase), *QDPR* (encodes dihydropteridine reductase), *NME2* (encodes nucleoside dephosphate kinase), and so on; the other included some genes encode regulatory proteins (e.g., *FTH1, GPX4*) and NADH dedydrogenase, which had stable mRNAs and unstable proteins. In addition, from the view of 3D structure, *NME2* is more stable than *GPX4* (**Figure S4**). Interestingly, there was no conserved fragment for gene sequences in cluster 1, which might be due to the existence of two sub-groups of genes in this cluster (**Figure S5**).

##### (ii) Cluster 2 and 3

There were 17 genes in each cluster, respectively. We have also attempted to interpret the combination of Cluster 2 and Cluster 3. The average global half-life of mRNAs and mRNAs in these clusters were not significantly different (*t-test p-values* were greater than 0.93 and 0.84, respectively) (**Figure 4c** and **Figure 4e**). On the other hand, these two clusters had different distribution in protein half-lives (**Figure 4d** and **Figure 4f**). Interestingly, there were conserved fragments for gene sequences in cluster 2 and cluster 3, respectively, while the conserved fragments for these two clusters were also similar to certain degree (**Figure S5**).

**Figure 4.**
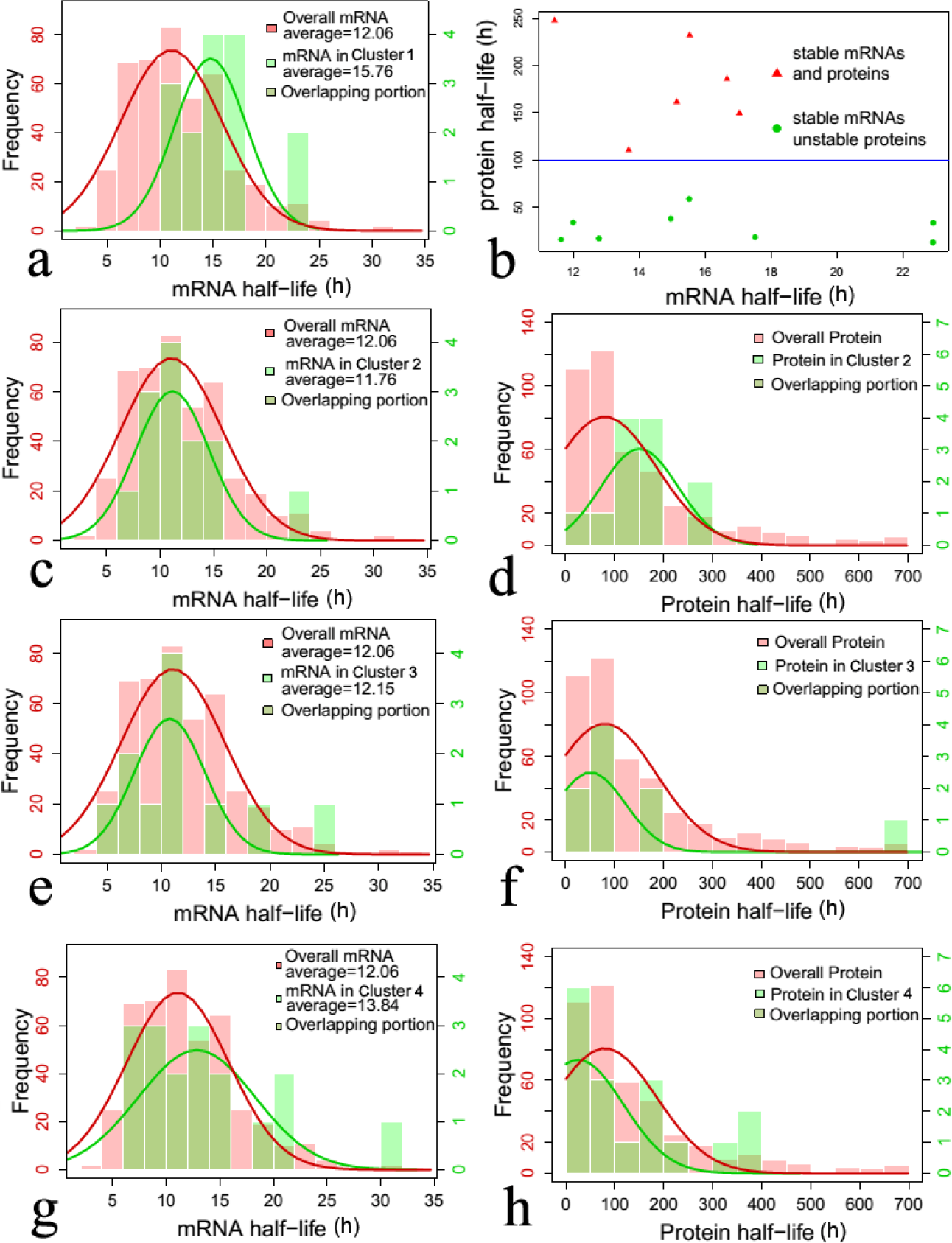
The mRNA and protein half-life histograms of clusters for mouse liver mitochondrial genes. The green bar refers to the secondary axis if the bar plot has one. (a) The histograms of the mRNA half-life distribution in Cluster 1 versus the global level distribution. (b) The mRNA and protein half-lives for Cluster 1 as shown in a scatter plot. (c) The histograms of the mRNA half-life distribution in Cluster 2. (d) The histograms of the protein half-life distribution in Cluster 2. (e) The histograms of the mRNA half-life distribution in Cluster 3. (f) The histograms of the protein half-life distribution in Custer 3. (g) The histograms of the mRNA half-life distribution in Cluster 4. (h) The histograms of the protein half-life distribution in Cluster 4. All half-lives were measured by hour as the basic unit.

##### (iii) Cluster 4

This cluster includes many ribosomal proteins. Compared with the half-lives of cytoplasmic ribosomal mRNA and protein, mitochondrial ribosomes had not only shorter-lived proteins (**Figure S6**), but also short-lived mRNAs. Interestingly, there was a conserved fragment for gene sequences in cluster 4 (**Figure S5**). These might be the reasons that 20 genes were clustered.

In summary, each of the clusters had specific half-life properties or protein functions, even gene sequence specificities, indicating that the bi-clustering results have uncovered the underlying dynamic relationship between gene expression and protein abundance.

### Feature Selection

Upon bi-clustering of the mouse liver mitochondria dataset, features that most appropriately describing and separating the clusters were identified. Different feature selection methods were applied, and their consensus results were used to build the prediction model.

According to cluster interpretation, we believed mRNA and protein half-lives could be most suitable for distinguishing genes from different clusters. Accordingly, we have analyzed 21 known features that are relatively well-studied, and have evidence to affect protein half-lives (**Table 1**). Other features associated with protein stability, such as the predicted instability index, were not included in this work but could be examined in future studies. Furthermore, 4 different methods: random forest, PCA and LDA analysis, PLS analysis, and mRMR were employed for feature selection analyses to identify the most important features (described in **Materials and Methods**).

Based on these consensus variables selected by multiple methods, we have finally selected 5 most representative features: C-secondary, Length, No._Lys_e, No._Lys and No._WCLTFYV (5 features would be enough to achieve the optimal balance between accuracy and efficacy (**Figure S7**)). (1) C-secondary represents the number of amino acids in secondary structure C (random coil), for which there are reports showing that the proportion of random coils in a protein structure has empirical relationships with protein stability^37^. (2) Length represents the protein sequence length, for which previous researches indicated that longer proteins tend to be more stable^31^. (3) No._Lys_e represents the number of exposed Lys, and (4) No._Lys represents the number of Lys in protein sequence. Lys residue has positive charge and very long flexible side chain, which prefers to be in protein exposed surfaces and interact with other partners. Furthermore, they are likely to perform as functional active sites of ubiquitin-protein ligases in the ubiquitin-proteasome system^34,38^, therefore Lys residues in proteins may affect the stability of the protein. (5) No._WCLTFYV represents the number of the amino acids W, L, C, T, F, Y in proteins. These 6 kinds of residues are large and hydrophobic which have also been shown in labile proteins by previous researches^31^. In summary, there are biological evidences to support that these 5 features are associated with protein stability. Therefore, we chose them for classification and building the prediction model.

### Comparison of prediction models

In this section, we have used the mouse liver mitochondrial gene expression and protein abundance data, and compared our novel CLM model with the previous reported two models: GLM and MARS. The GLM model calculates a single linear relationship between mRNA and protein abundances for all genes. The MARS Model performs prediction based on subsets of genes (separated by segmentation based on all genes). MARS differs from the CLM model in that the subsets of genes are selected based on automatic selection that would yield best adaptation of linear model to datasets, while in CLM model the subsets are selected by bi-clustering.

#### (1) Comparison with the GLM model

For GLM model, the average prediction accuracy of 27.0% was achieved, using 100 genes as the training dataset and the last (342 genes) as the testing dataset (input: the mouse liver mitochondrial dataset). For CLM model, the genes as training and testing datasets were carefully selected basing on clustering results from bi-clustering, which would ensure the accuracy and fairness of the prediction model evaluation. One linear model was built for every cluster obtained from bi-clustering. Through this process, the average accuracy based on 7 clusters was 35.09%, which was larger than that in GLM model on total genes (27%). Although the number of genes in these clusters (123 genes) were relatively small, they were representative enough for the general patterns that connect gene expressions and protein abundances.

In addition, we discovered several genes that can only be predicted accurately by CLM model (the amaranth triangles in black circle in Figure 5), e.g., *Nags* and *Surf1.* The *Nags* gene encodes malonyl N-acetylglutamate synthase that catalyzes the production of N-acetylglutamate from glutamate and acetyl-CoA. This enzyme is important for mammals, because it produces the regulator of urea cycle N-acetylglutamate, which activates carbamoyl phosphate synthetase I, catalyzing the initial reactions of urea cycle. Since the gene expression and protein abundance are relatively low, the linear model on total genes cannot predict this gene. The same reason applies to *Surf1* gene. *Surf1* gene encodes surfeit gene 1 that may play both a positive and negative regulatory role in gene expression^39^. The mRNA and protein half-lives of the *Surf1* gene were 10.61 hour and 36.03 hour, respectively, which is shorter to reflect its regulatory function, compared with the average mRNA and protein half-lives. These cases indicated the advantage of CLM model’s divide-and-conquer approach over that of GLM.

**Figure 5.**
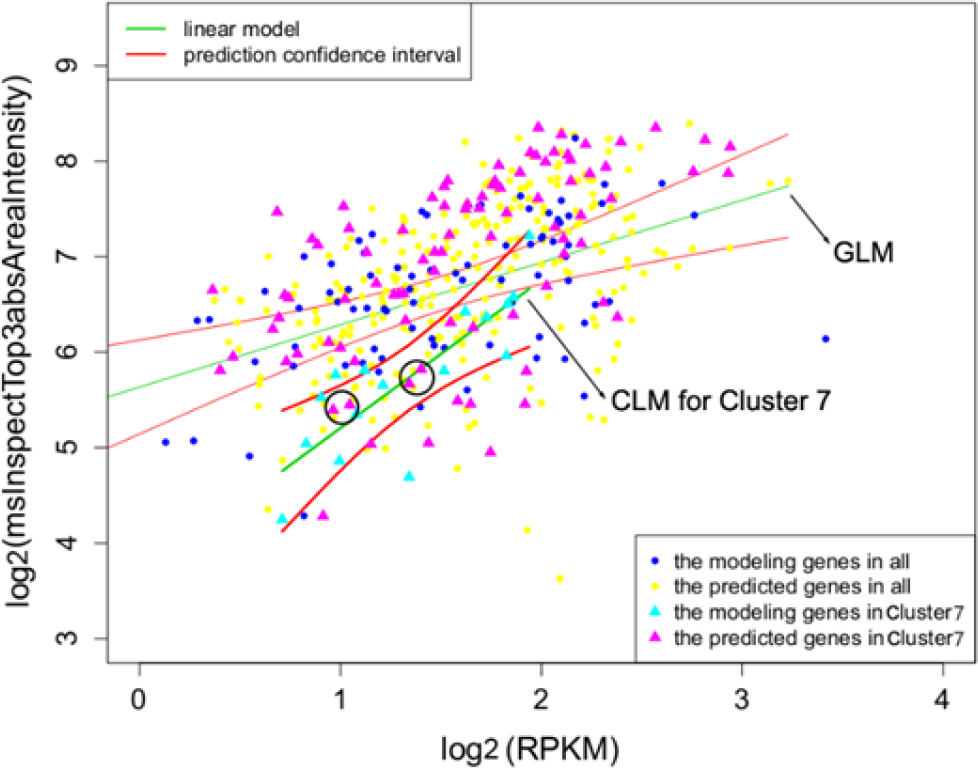
Results of GLM and CLM modeling for mitochondrial genes from mouse liver tissue. The amaranth triangles in black circle are representatives of genes that can only be predicted accurately by CLM model for different clusters. Notice that in CLM model for Cluster 7, gene *Nags* is in the left black circle and gene *Surf1* is in the right black circle.

#### (2) Comparison with the MARS model

In the previous study on MARS^21^, the correlation between real values and predicted values of protein abundance was considered as measure of model quality. However, “good correlation” is not equivalent to “accurate prediction”. Error Sum of Squares (SSE) between real and predicted values was another quality metric to compare CLM and MARS model.

Based on the above 5 consensus features (C-secondary, Length, No._Lys_e, No._Lys and No._Lys_e/L), CLM model could obtain better prediction results, with higher Pearson correlation (r) and lower SSE between real values and predicted values in 3 clusters (Cluster 2, 3 and 6) (**Table S2**). *i)* In Cluster 2 and 3, we observed that the correlation coefficients between mRNA abundance and protein abundance was high (0.91 and 0.88, respectively) and genes in these two clusters deviated from the global distribution (**Figure 3**). In addition, the genes of the two clusters had similar average half-life of mRNAs compared with that of global mRNAs (*t-test p-values* were greater than 0.93 and 0.84, respectively), yet different distributions in protein half-lives (**Figure 4**). *ii)* In cluster 6, genes did not have high correlation and deviated from the global distribution, the properties of these genes were similar to those in Cluster 3 that included some ribosomal proteins. And there was no significant difference between global mRNA half-lives and those in this cluster (*t-test p-values* was 0.54). We concluded that CLM model might have superior prediction power for clusters whose mRNA half-lives were not significantly different from global mRNA half-lives. We conjectured that CLM was better than MARS for these clusters, mainly because for each cluster, predictions based on CLM were more biologically meaningful as regard to genes within the clusters, while predictions based on MARS might be affected by genes outside of the cluster but in the same segment based on which the spline was generated.

### Validation of the prediction method: Applicability of computational model on other samples

Due to the existence of various transcriptome datasets, the features for protein prediction might be diverse, thus the features selected above would be inconsistent among different datasets. Rather, the entire procedure to select features as well as building the prediction model should be generally applicable and superior to simpler models like GLM model. Such a proof can only be given with validation based on different datasets. Therefore, we have validated the entire prediction procedure based on three additional datasets: mitochondrial genes from the mouse brainstem tissue (with relatively more stable turnover), genes in *Saccharomyces cerevisiae* (with relatively more dynamic turnover), as well as genes in embryonic development process in *Danio rerio* (with relatively more dynamic turnover).

#### (1) Mitochondrial genes from mouse brainstem tissue

Similar to mitochondrial genes from mouse liver tissue, the data of transcripts, proteins and associated quantitative information in mouse brainstem tissue were obtained from the same work as used in Ning *et al.*^11^. We followed the same analysis procedure of bi-clustering, feature selection and prediction model building as above, in which the “prediction model building” module was repeated many times to reduce randomness in classification. The average accuracy of multiple clusters was 28.87% based on 5 features (E-secondary, No._WCLTFYV, No._Lys, No._Lys_e/L and No._Lys_e). This accuracy was higher than that from all-gene-based linear correlation of 23.37%. When compared with MARS Model, CLM model could have a better performance (**Table S3**).

Some interesting prediction results, such as those related with half-lives and protein functions, have also been observed. *i)* Gene *Supv3l1* encodes ATP-dependent RNA helicase. The mRNA and protein half-lives of the gene *Supv3l1* were 6.88 hour and 51.96 hour, respectively. Both the mRNA half-life and the protein half-life were shorter, compared with the overall average. This might be because the helicase only takes part in the initiation of material replication process which is transient and happens occasionally. Therefore, short half-lives for mRNA and protein reflect the principle of the adaptation of the properties and function. In addition, the reason that the GLM model on all genes cannot predict the protein abundance of gene *Supv3l1* might be that mRNA and protein abundance were relatively low. *ii)* The same reason applied to gene *Ppif.* The *Ppif* gene encodes Polymerase delta-interacting protein. The mRNA and protein half-lives of the gene *Ppif* were 5.49 hour and 160.09 hour, respectively. The mRNA half-life is shorter, and the protein half-life is longer compared with the overall average. Although this protein participates in genetic material replication process, it is indispensable to function for a relatively long time, while the corresponding mRNA will degrade immediately after translation. These two genes’ protein abundance could not be predicted accurately by GLM model, but CLM model could predict it with good accuracy (**Figure 6**).

**Figure 6.**
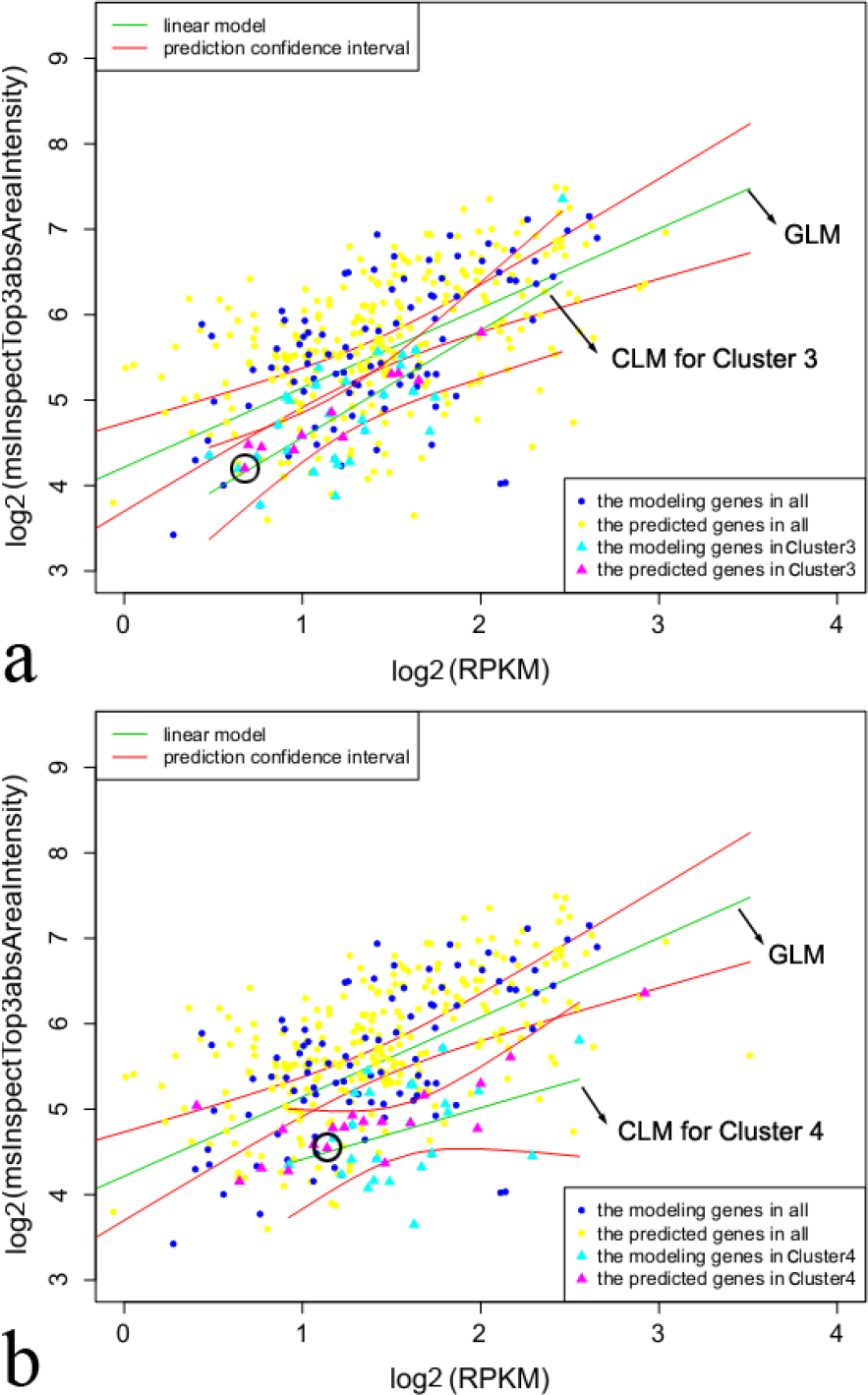
Results of GLM and CLM modeling for mitochondrial genes from mouse brainstem tissue. (a) GLM model and CLM model for Cluster 3 and gene *Supv3l1* in circle. (b) GLM model and CLM model for Cluster 4 and gene *Ppif* in circle. The amaranth triangles in black circle are representatives of genes that can only be predicted accurately by CLM model for different clusters.

#### (2) Genes from Saccharomyces cerevisiae

The transcripts, proteins and associated quantitative information for *Saccharomyces cerevisiae* were obtained from Griffin *et al.*^25^. After feature selection, the most important variables selected include Length (L), No._WCLTFYV, No._EDKNRQ, as well as No._Lys_e/L, No._Lys_e, No._Lys, H-secondary, C-secondary, PEST/L. All 5 variables selected from the “mouse liver mitochondrial” dataset were also deemed most important for *S. cerevisiae.* But a couple of variables (e.g., No._PEST and Exposed%), which were not considered important for the “mouse liver mitochondrial” dataset, were important for discriminating different genes in *S. cerevisiae* samples. Based on this different set of selected features (again the top 5 features that include C-secondary, No._Lys_e, No._Lys, No._Lys_e/L, No._PEST), a new prediction model was built. Results have shown that the prediction accuracy (49.4%) could be higher than that from all-gene-based linear correlation (42.4%), and also higher than directly using the prediction model based on the “mouse liver mitochondrial” dataset (45%).

#### (3) Genes from Danio rerio

To further explore the applicability of our method, datasets including transcripts, proteins and associated quantitative information for *Danio rerio* in embryonic development process were obtained from Shaik *et al*^26^. After feature selection, the most important variables selected include Length, Exposed%, NO._WCLTFYV, NO._EDKNRQ, and EDKNRQ\L. By comparing with the results for the other three datasets, we discovered the variable, NO._WCLTFYV, is a relatively more universal and important feature in the correlation analysis of mRNA and protein for these four the datasets/species. Meanwhile, the variable EDKNRQ\L, which was not considered important for either two “mouse” datasets or *S. cerevisiae* dataset, was important for discriminating different genes in the embryonic development process of *Danio rerio*. It was discovered previously that the above charged amino acids (E, D, K, N, R, Q) were enriched in more stable proteins^33^. Thus, the variable of EDKNRQ\L might be much more related to protein stability or the process of selective protein degradation (mostly ubiquitin-mediated)^38^, which might be more critical for differentiation/signal transduction in the process of embryonic development in *Danio rerio* dataset than in the other three datasets.

In summary, when the other three independent datasets have been used in this study, the prediction accuracy of CLM model could reach a higher accuracy compared to MARS model. Meanwhile, in *Danio rerio* dataset, the comparison results between MARS Model and CLM model demonstrated that CLM performed better than MARS, especially in Cluster 1 (MARS: R^2^=0.37, SSE=17200; CLM: R^2^=0.48, SSE=12720) and Cluster 3 (MARS: R^2^=0.45, SSE=2550; CLM: R^2^=0.52, SSE=2326), for which higher correlation coefficient values and smaller SSE values could be observed from CLM model (**Table S3**).

## Conclusion

Identification of major determinants for the correlation between gene expression and protein abundance can lead to better understanding of gene regulation mechanisms and better prediction of protein abundance from gene expression data. In this work, we have proposed a bi-clustering method to cluster genes that have consistent patterns for the correlation between gene expression and protein abundance. The clustering results have been interpreted by the properties of both transcripts and proteins, which showed that mRNA half-life, protein half-life and protein 3D structure could affect gene expression and protein abundance profoundly. Based on these results, we have proposed a CLM model for prediction of protein abundance based on gene expression and protein features.

On the mouse liver and brainstem mitochondrial datasets, this approach worked well for protein abundance prediction from gene expression data, proving the validity of the prediction model on mouse mitochondrial genes. Additionally, on *Saccharomyces cerevisiae* transcriptomic and proteomic data, the CLM model was built on a different set of features, and the prediction accuracy could again reach a satisfactory level better than all-gene-based GLM model. Furthermore, relatively satisfactory results of this CLM modeling approach were also observed based on *Danio rerio* embryonic development dataset. Based on these results for four datasets representing diverse kinds of species and organelles, it was quite clear that for most clusters, CLM models could achieve higher correlation and lower SSE between real values and predicted values of protein abundance compared to MARS models.

Therefore, we conclude that the general approach to select features as well as building the CLM prediction model can achieve relatively high accuracy for a wide range of datasets. Furthermore, the 5 most important protein features differ among species and tissues, resulting in different CLM models, and reflect different mRNA and protein turnover for diverse kinds of species and organelles.

We also noticed that the prediction accuracies for all mouse genes/proteins were still far from perfect and quite low when using the prediction model built based on the “mouse liver mitochondrial” dataset. This may reflect a specificity of each dataset, indicating that there is still room for improving prediction model. And the improved prediction model might be built based on further development of optimization methods for feature selection, as well as non-linear model tuning. All these might help for better prediction of protein abundances from gene expressions without much more other information.

## Availability

The computational analysis pipeline for CLM methods, as well as manuals and example datasets are provided online at: http://www.microbioinformatics.org/software/clm.html.

## Authors’ contributions

XJW, LT and KN designed the whole framework, analyzed the data and wrote the initial draft of the manuscript. HCZ, DBB and XQG provided the results of protein 3D structure. XJW, QZ, XQS, XPC, XQG, AP and KN reviewed and revised the manuscript.

## Acknowledgments

The study was funded by the Ministry of Science and Technology’s National Basic Research Program of China (973 Program) grant 2012CB316502, the National Nature Science Foundation of China under grants 61103167, 31271410, 11175224, 11121403, 31270834 and 61272318, the Open Project Program of State Key Laboratory of Theoretical Physics (No. Y4KF171CJ1). This work made use of the Infrastructure provided by the European Commission co-funded project CHAIN-REDS (GA NO. 306819). We would also thank Dr. Shiwei Sun and Dr. Francis Ng for discussions about building prediction model, as well as for careful proof-reading.

